# Role of inhibitory control in modulating focal seizure spread

**DOI:** 10.1101/146407

**Authors:** Jyun-you Liou, Hongtao Ma, Michael Wenzel, Mingrui Zhao, Eliza Baird-Daniel, Elliot H Smith, Andy Daniel, Ronald Emerson, Rafael Yuste, Theodore H Schwartz, Catherine A Schevon

**Affiliations:** Department of Physiology and Cellular Biophysics, Columbia University, New York, NY 10032, USA; Department of Neurological Surgery, Feil Family Brain and Mind Research Institute, Sackler Brain and Spine Institute, Weill Cornell Medical College, New York, NY 10065, USA; Neurotechnology Center, Department of Biological Sciences, Columbia University, New York, NY 10027, USA; Department of Neuroscience, Columbia University, New York, NY 10032, USA.; Department of Neurological Surgery, Columbia University Medical Center, New York, NY, 10032, USA.; Hospital for Special Surgery, Weill Cornell Medical College, 535 East 70th Street, New York, NY 10021, USA.; Department of Neurology, Columbia University Medical Center, New York, New York 10032, USA.

**Keywords:** Acute seizure model, inhibitory restraint, 4-aminopyridine, neocortical seizure model, epileptic network

## Abstract

Focal seizure propagation is classically thought to be spatially contiguous. However, distribution of seizures through a large-scale epileptic network has been theorized. Here, we used a multielectrode array, wide field calcium imaging, and two-photon calcium imaging to study focal seizure propagation pathways in an acute rodent neocortical 4-aminopyridine model. Although ictal neuronal bursts did not propagate beyond a 2-3 mm region, they were associated with hemisphere-wide field potential fluctuations and parvalbumin-positive interneuron activity outside the seizure focus. While bicuculline surface application enhanced contiguous seizure propagation, focal bicuculline microinjection at sites distant to the 4-aminopyridine focus resulted in epileptic network formation with maximal activity at the two foci. Our study suggests that both classical and epileptic network propagation can arise from localized inhibition defects, and that the network appearance can arise in the context of normal brain structure without requirement for pathological connectivity changes between sites.

## Introduction

The topology of seizure propagation has long been debated, and may depend on factors such as cortical connectivity and underlying pathology. Classically, seizures are thought to spread to physically contiguous regions, expanding in a sequential manner through progressive recruitment of adjacent cortical circuitry (Supplementary Figure S1A). This progression manifests clinically during the well-known Jacksonian march (Extercatte *et al*., 2015), and has been extensively documented in animal models, both in vivo and in vitro (Bikson *et al*., 2003; Pinto *et al*., 2005; Trevelyan *et al*., 2006; Trevelyan *et al*., 2007b; Wenzel *et al*., 2017), and in spontaneous human seizures (Schevon *et al*., 2012; Smith *et al*., 2016). However in the clinical setting, many seizures appear to spread rapidly through large brain areas, leading to the widely held notion that large-scale network behavior plays a causative role(Kramer *et al*., 2010; Luo *et al*., 2014; Khambhati *et al*., 2016) (Supplementary Figure S1B). Furthermore, recruitment of distant sites cannot be explained by the slow pace of contiguous seizure spread (< 1 mm/sec) and relatively short seizure durations.

Epileptiform discharges during focal seizures can be observed in EEG recordings at a significantly larger spatial scale than its neuronal substrates. Analyses of in vivo human microelectrode recordings suggests that widespread synaptic barrages emanating from a small ictal core are responsible for the visual EEG appearance of far-ranging seizure spread (Schevon *et al*., 2012; Weiss *et al*., 2013). The resulting spatial discrepancy between the ictal core, i.e. seizing brain territory in which neurons demonstrate pathologically intense, time-locked, synchronized bursts, and the region over which epileptiform field potential activity may be detected, has confounded previous efforts to investigate long-range seizure spread, both in animal models and in human epilepsy. Thus, new investigations into long-range spread mechanisms are warranted.

Here, we propose a scenario that can explain the emergence of large-scale network activity via compromise of surround inhibition at distant sites, resulting in secondarily activated seizure foci. Detailed in vitro brain slice studies have shown that in normal tissue, a fast, strong inhibitory conductance in adjacent cortical territory prevents the distributed synaptic activity from triggering ictal activity (Trevelyan *et al*., 2006; Trevelyan *et al*., 2007b), and evidence of the same process has been detected in humans (Schevon *et al*., 2012; Eissa *et al*., 2017). To date, the investigations of surround inhibition have been largely limited to regions immediately adjacent to the invading seizure. In our scenario, surround inhibition not only plays an important role in containing seizure invasion to adjacent cortex, but also protects areas distant to the recruited region. We hypothesize that strongly excitatory synaptic projections arise from the seizure focus and are distributed widely, but are normally masked by local inhibitory responses.

We assess the effects of ictal activity projection at varying cortical distances, as well as inhibitory responses using multielectrode arrays, wide field calcium imaging, and two-photon calcium imaging of interneurons in the acute rodent in vivo 4-AP model. The in vivo 4-aminopyridine (4-AP) seizure model replicates the key features of propagating seizures, including characteristic epileptiform discharges and localized neurovascular coupling effects at the application site (Bahar *et al*., 2006; Ma *et al*., 2009; Zhao *et al*., 2009; Zhao *et al*., 2011; Ma *et al*., 2013). We then introduce bicuculline methiodide (BMI) to block GABA-A activity at varying distances from the 4-AP focus, to simulate the effects of different topologies of locally impaired inhibition. The purpose of our dual-site model is to demonstrate that seizure propagation can proceed via two topologically distinct patterns that both depend on the same inhibition breakdown process: one involving slow (< 1 mm/sec), contiguous spread and the other operating across large distances at much greater speeds. This provides a plausible mechanism reconciling the longstanding, classical contiguous seizure propagation patterns with the observed large-scale epileptic network behavior during human seizures (Smith and Schevon, 2016).

## Materials and Methods

A total of 20 animals were used for the three experimental modalities in this study: multielectrode array recording (8 animals), wide-field calcium imaging (10 animals), and two-photon calcium imaging (2 animals). All experimental procedures were approved by either the Weill Cornell Medical College or Columbia University Animal Care and Use Committee following the National Institutes of Health guidelines.

### Multielectrode array recording

#### Animal preparation

Adult male Sprague–Dawley rats (200–350 g) were anesthetized with isoflurane in 70% N_2_:30% O_2_, 4% induction, and 1.5–2% maintenance. Body temperature was maintained at 37 °C with a regulated heating blanket (Harvard Apparatus, Holliston, MA). The heart rate, pO_2_, and the end tidal carbon dioxide (EtCO_2_) were carefully monitored with a small animal capnography (Surgivet, Waukesha, WI) and were sustained throughout the experiment (heart rate: 250–300 pulse/minute, pO2 > 90%, EtCO_2_ ~25–28 mmHg). The exposed brain was covered with cotton balls socked with artificial cerebrospinal fluid (ACSF) to preserve cortical moisture.

#### Array configuration

Two types of multielectrode arrays were used in this study. The first type, used for unilateral (left) hemisphere recording, was configured into a 10 by 10 grid with 400 µm interelectrode distance (Blackrock Microsystems Inc, Salt Lake City, UT). The second type, used for bilateral hemisphere recording, was configured into two separate 10 by 5 grids with 400 µm interelectrode distance. Both types of arrays were implanted into the cortex with a 0.5 mm pneumatic implanter (Blackrock Microsystems Inc, Salt Lake City, UT)(Rousche and Normann, 1992). The grids’ anterior edges met the animal’s somatosensory cortices. In the bilateral array experiment, the long side (10 electrodes) was aligned anterior-posteriorly, and each grid was used for each hemisphere. Reference electrodes were placed subdurally, distal to recording sites, and dynamically selected to minimize reference artifacts.

#### Array Signal recording

Raw electrical signal was digitally sampled at 30 kHz with 16-bit precision (range ±8 mV, resolution 0.25 *µ*V, 0.3 to 7500 Hz band pass). Raw data were subsequently separated into two frequency bands: multiunit activity (MUA, 300 to 3000 Hz, 512^th^ order zero-phase shift, window-based FIR filter). LFP was derived by downsampling the raw signal to 1 k Hz after anti-alias filtering (500 Hz low-pass, 90^th^ order, zero-phase shift, window-based FIR filter). Channels with background noise amplitude (estimated by scaled median absolute deviation) more than 8 μV in MUA band or excessive paroxysmal artifacts were discarded (Quiroga *et al*., 2004).

#### Multiunit spike detection & firing rate estimation

Multiunit spikes were detected from the MUA band by a threshold crossing method(Quiroga *et al*., 2004). Each channel’s detection threshold was set to be −5 s.d. of the channel’s background noise (estimated from a 2-minute baseline recording). Detection refractory period was set to be 1 ms, to minimize detection of multiple peaks due to noise. Channels that failed to detect more than 1 multiunit spike per minute were also excluded. For each channel, instantaneous multiunit firing rates were estimated by convolving its spike train with Gaussian kernels, sampled every 1 ms (Bokil *et al*., 2010; Shimazaki and Shinomoto, 2010; Smith *et al*., 2016). Two types of Gaussian kernels were used in this study: a 10-ms s.d. Gaussian kernel for capturing rapid change of multiunit firing rates and a 1-second s.d. kernel for slow change of average firing rates.

#### Seizure model

4-Aminopyridine (4-AP, Sigma-Aldrich, 15 mM, 500 nL), a potassium channel blocker, was injected at the anterior edge of the array in the left somatosensory cortex, targeting 300–500 μm below the cortical surface through a glass microelectrode using a Nanoject II injector (Drummond Scientific, Broomall, PA) (Pongracz and Szente, 1979; Rutecki *et al*., 1987; Avoli *et al*., 2016; de Curtis and Avoli, 2016) after a 2-minute baseline recording. The 4-AP dose was increased to 1000 nL if electrographic seizures were not observed in the next 20 minutes of recording. 30 to 60 minutes after the first electrographic seizure, bicuculline methiodide (BMI, Sigma-Aldrich, 5 mM), a GABA-A receptor antagonist, was introduced either by bath application (left hemisphere) or focal injection (500nL, 300–500 μm depth at left visual cortex or right somatosensory cortex). The BMI injection site was placed at least 4 mm distant from the 4-AP site, in order to ensure that the region affected by each agent (2mm diameter, Schwartz and Bonhoeffer, 2001) would not overlap. Neural dynamics were recorded for another 30 to 60 minutes following BMI injection.

#### Ictal activity detection and quantification

The line-length feature of LFP was used to quantify ictal activity and facilitate systematic and objective ictal event detection (Esteller *et al*., 2001; Guo *et al*., 2010). For each channel, line-length was calculated for 1-second moving windows (step size: 1 ms). Periods that had more than 3 channels demonstrating greater than 3 times more line-length than the baseline and lasted for more than 5 seconds were considered as ictal event candidates, which were further visually reviewed to exclude artifacts. The onset and offset of ictal events were adjusted manually to encompass all epileptiform discharges. Furthermore, two ictal events were required to be at least 20 seconds apart temporally to be considered two isolated episodes (Ma *et al*., 2009; Zhao *et al*., 2009; Zhao *et al*., 2011). The periods between two consecutive ictal events are defined as interictal periods. For each ictal event, the ictal center is defined as physical location of the electrode where we detected the highest line-length.

#### Multiunit spike-LFP coupling

We used spike-triggered averaging to investigate the association between multiunit firing and LFP dynamics of the entire region sampled by microelectrodes (Schwartz *et al*., 2006; Eissa *et al*., 2016). Multiunit spikes which were detected within 1 mm from the ictal center were combined to create the spike train used for averaging.

#### Cross-correlation of multiunit spike trains

Cross-correlation between binned spike-trains (1 ms resolution) was used to determine the temporal relationships between any two spike trains. The timing of peak cross-correlation,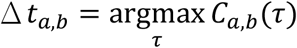 was defined as the temporal delay between two channels, *a* and *b*.

#### Cluster analysis of multiunit spike trains

Non-negative matrix factorization was applied to study the clustering phenomenon of ictal dynamics (Lee and Seung, 1999; Hutchins *et al*., 2008). During each ictal event, each channel’s instantaneous multiunit firing rate was determined by convolution with a 10-ms s.d. Gaussian kernel. The resultant time series were sampled every 1 ms, aligned in rows, and assembled into a data matrix, *A*, where *A*_*i*_,_*j*_ is the instantaneous firing rate of channel *i* at time *j*. The matrix, *A*, consisting of all non-negative entries, was further factorized into two non-negative matrices, *A* ≈ *WH* with latent dimensionality *k* = 2.

## Wide-field Calcium Imaging

### Calcium dye loading and fluorescence measurement

The calcium indicator Oregon Green 488 BAPTA-1 AM (OGB-1, Life Technologies, Grand Island, New York) was employed for recording of neural activity. Convection enhanced delivery (CED) was employed to bulk load the entire neocortex with OGB-1(Ma *et al*., 2014b). In brief, 50 µg of OGB-1 was diluted in 5 µL of DMSO-F127 then in 50 µL of ACSF. 8 µL of OGB-1 solution was injected in the neocortex, via a glass electrode (50-100 µm opening) placed ~1 mm below the brain surface, at the speed of 100 nL/min, using a micro-pump (WPI, Sarasota, Florida). A CCD camera (Dalsa camera in Imager 3001, Optical Imaging, Rehovot, Israel) using a tandem lens (50 × 50 mm) arrangement was focused 300–400 μm below the cortical surface. A 470±10 nm LED was employed as the illumination source for calcium-sensitive dye and illumination was directed using fiber-optic light guides. A 510 nm long-pass filter was placed before the camera to prevent calcium illumination contamination, while permitting calcium dye signal.

### Calcium imaging analysis

Data were analyzed by customized MATLAB functions. Calcium images were convolved with a spatial Gaussian kernel (σ = 3 pixels) to increase signal-to-noise ratio. The signal changes were calculated as dF/F, where F was the baseline illumination when no epileptiform discharges were noticed and dF was the signal change during epileptiform activity as identified by the LFP. A 1 Hz high-pass filter was applied to the calcium imaging data to remove the calcium signal from glial cells (Ma *et al*., 2014a).

A seed trace initiated correlation method(White *et al*., 2011) was employed to calculate the spatial spread of neuronal activity. Briefly, a seed trace was selected from a small region of interest (ROI) from the 4-AP and BMI injection sites. The correlation coefficients between the seed trace and trace from every individual pixel in the surrounding image were calculated. A heat map was generated using the correlation coefficient (CC) at each pixel.

### Two-photon Calcium Imaging

#### Animal preparation, seizure model, and ictal event detection

PV-Cre mice (RRID:IMSR_JAX:017320) were crossed with LSL-GCaMP6F(Chen *et al*., 2013) mice (RRID:IMSR_JAX:024105), resulting in GCaMP6F expression specifically in parvalbumin (PV) positive interneurons. Animals were anesthetized and their physiological conditions were maintained similarly to protocols described in previous sections. Two craniotomies over the left hemisphere were created – one (posterior primary visual cortex) for 4-AP (15 mM, 500 nL) injection and the other (primary somatosensory cortex just posterior to bregma, ~4 mm anterior to 4-AP injection site) for imaging.

#### Image collection

Calcium imaging of GCaMP6F positive interneurons was performed using a two-photon microscope (Bruker; Billerica, MA) and a Ti:Sapphire laser (Chameleon Ultra II; Coherent) using a 25x objective (water immersion, N.A. 1.05, Olympus). GCaMP6F was excited with a laser wavelength of 940 nm, fluorescence emission was collected through a 535nm (green) filter (Chroma, Bellows Falls, VT). Resonant galvanometer scanning and image acquisition (frame rate 30,206 fps, 512 × 512 pixels, 150-200 µm below the pial surface) were controlled by Prairie View Imaging software.

#### Image analysis

Regions of interest (ROIs) were registered manually to target GCaMP6F-expressing PV(+) interneurons using ImageJ (National Institute of Mental Health, Bethesda, MD.). To minimize cell signal contamination by surround neuropil fluorescence changes, we applied ROI shrinkage (Radial subtraction of 1 pixel from somatic ROI)(Hofer *et al*., 2011). Individual cell fluorescence was calculated as the average across all pixels within the ROI. Then, we calculated relative changes in fluorescence (F) as 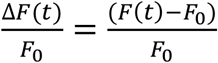 where *F*_0_ represents the mean of the lowest 25% of values within a 500-frame window centered around each value F(t). Finally, to minimize tissue pulsation artifacts in the extracted traces, individual traces were filtered with a 2-second LOWESS smoothing envelope.

#### Ictal events – PV interneuron activity coupling

LFP, recorded from a glass micropipette (2-5 MΩ, AgCl wire in isotonic PBS solution) 100 µm beneath the pial surface near the 4-AP injection site in primary visual cortex (V1), were used for ictal event detection. A reference electrode was positioned over the contralateral frontal cortex. LFP signals were amplified by a Multiclamp 700B amplifier (Axon Instruments, Sunnyvale, CA), low-pass filtered (300 Hz, Multiclamp 700B commander software, Axon Instruments), digitized at 1000 Hz (Bruker) and recorded using Prairie View Voltage Recording Software along with calcium imaging. Ictal events were inspected visually. Specific ictal onset time was determined by searching the turning point of initial negative DC-shift: 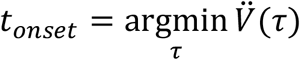

To calculate time lags between individual ictal onsets (measured by LFP within the seizure initiation site) and the respective time of local PV population recruitment at the FOV in primary somatosensory cortex (distance: ca. 4 mm anterior to initiation site), we derived an average calcium transient of all imaged PVs in the FOV. Then, we defined timing of maximum curvature (most negative 2^nd^ derivative) of the population calcium transient during each peri-ictal onset window (centered around the time point of the electrographic seizure onset) as the time point of PV population recruitment. Eventually, the lag was calculated as Δ*t* _*seizure* =_ *t_calcium_ –t_LFP_*.

## Results

### Focal 4-AP injection induced spatially constrained ictal events

Figure 1A shows the experimental setup. Baseline recordings were taken to rule out the presence of pre-existing epileptiform activities prior to pharmacological manipulation. Injecting 4-AP induced focal seizure-like events, which we refer to as ictal events for simplicity of presentation (Figure 1B-C). These resembled low-voltage fast onset electrographic seizures that are commonly seen in humans (Lee *et al*., 2000; Perucca *et al*., 2014). In some cases, twitching of the contralateral or bilateral limbs and whiskers was noted, temporally correlating with the epileptiform discharges. Epileptiform discharges (Figure 1B, anterior-right corner of the microelectrode array) and synchronized multiunit bursts (Figure 1C) were detected at the 4AP injection site. During ictal events, similar to ictal cores of human focal seizures (Merricks *et al*., 2015), the waveforms of multiunit spikes near the ictal center were distorted (Figure 1C, light blue arrows), whereas multiunit waveforms were preserved at distal sites (Figure 1C, yellow arrows). The spatial extension of the epileptiform LFP activities remained stable during repetitive ictal events (Figure 1D). Overall, both LFP (local field potential) activity and multiunit firing rates decreased with increasing distance from the 4-AP injection site in all experiments, with MUA (multiunit activity) showing more rapid decline. Increased LFP line length was detectable beyond the region defined by MUA firing greater than baseline (Figure 1E-F, see legends for detailed statistics).

**Figure 1.**
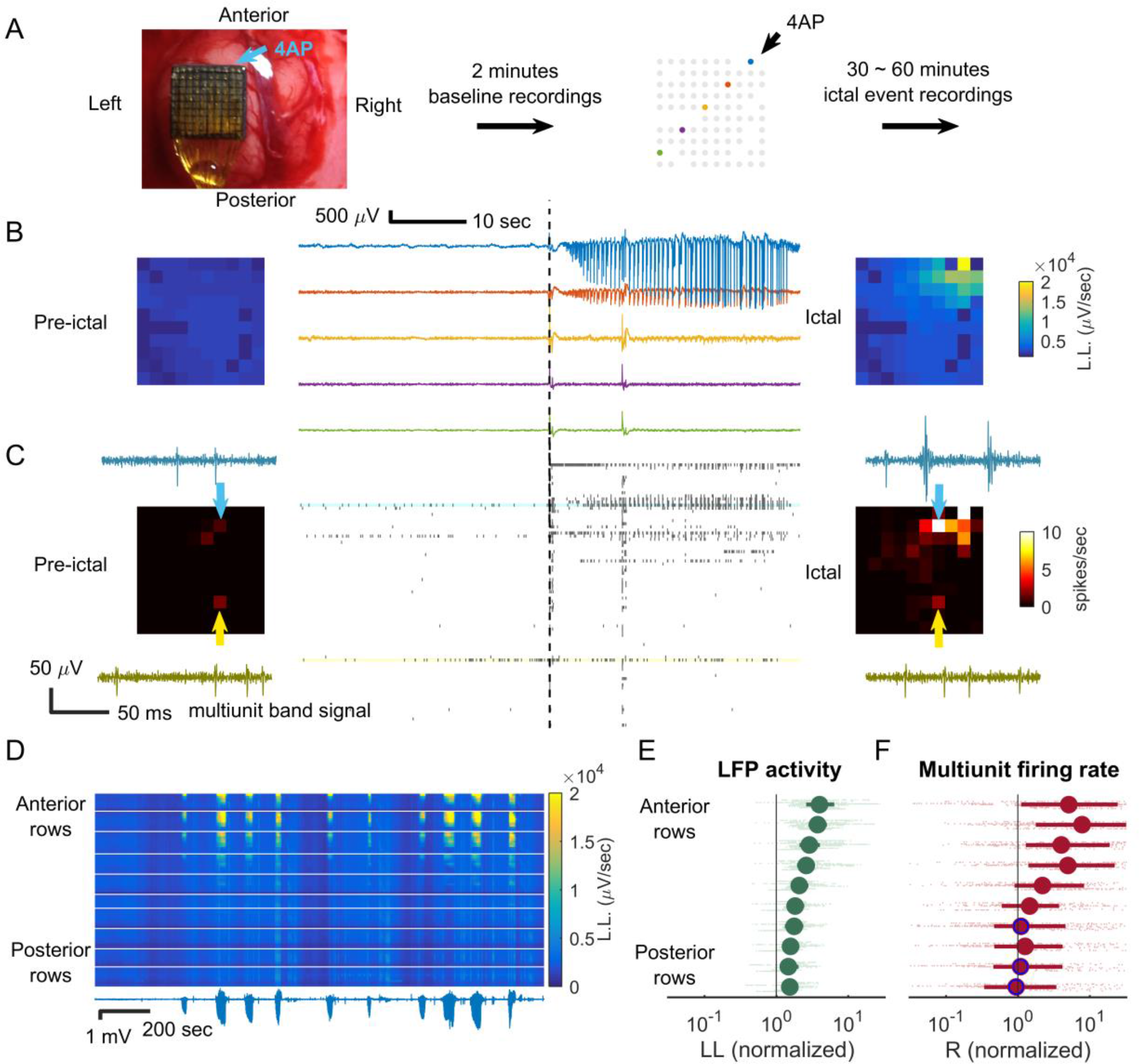
Focality of 4-AP ictal events recorded with microelectrode array. A. Experimental setup. The microelectrode array was implanted in the left hemisphere. 4-AP was the injected at the anterior border (upper right corner) to induce ictal events. Missing dots denote non-recording electrode sites. Five microelectrodes are highlighted with a color code corresponding to traces shown in ensuing panels. B. Sample LFP of pre-ictal (left) and ictal (right) periods after 4AP injection. Line length (LL) is used in corresponding color spectrum plots to show the spatial layout of ictal activity. Note the focality of the event. C. MUA raster plot during pre-ictal (left) and ictal (right) periods. The time axis (horizontal) aligns with Panel B. Average multiunit firing rates are shown in the corresponding heat maps. The multiunit band signal of sample channels (light blue and yellow) is shown in the subpanels. The spatial extent of the activity is more restricted than is the case for LFP. D. Repetitive ictal events from one animal are shown. Ictal activities were similarly spatially distributed - concentrated at the anterior region sampled by the multielectrode array. Lower trace: LFP recording from the ictal center (highest line length). The corresponding color map shows the activity of each channel over time. Displayed from top to bottom, channels are arranged by their location in the microelectrode grid, grouped according to row with white dividing lines between rows, and arranged right to left within each row. Activity is denoted by line-length averaged over 5 second moving windows. E. Quantification of ictal activity along the anterior-posterior axis. Average line-length of each channel was calculated for all channels and each ictal event, normalized to the baseline (8 animals, 120 events, data points in light green). The vertical axis is arranged by channel row, with each animal’s data represented with minor vertical offsets. Green markers and error bars represent the row-specific median and interquartile ranges. LFP activity increased in all rows (sign test, p<0.001). There was a clear anterior-posterior gradient (Spearman’s correlation coefficient, row position vs. line-length: 0.59, 8909 data points, p << 0.001). F. Quantification of multiunit firing rate along the anterior-posterior axis for the data set of Panel E, with the same presentation conventions. The anterior channels, closest to the 4-AP microinjection site, showed a significant multiunit firing rate increase (sign test, top 6 rows, p<0.001) but not the posterior channels (3 of bottom 4 rows, markers with blue rims, p=0.26, 0.07, and 0.53 from anterior to posterior respectively). Spearman’s correlation coefficient again confirmed the existence of an anterior-posterior gradient (*ρ* =0.31, 3915 data points, p<<0.001). Note the sharp drop in firing rates in the transition from rows 4 to 6, in contrast to the smooth gradient seen in LFP activity (Panel E).

### Wide-field calcium imaging confirmed 4AP ictal event’s focal nature is regionally invariant

To overcome the limited spatial sampling of the microelectrode array (4 mm × 4 mm), we used wide-field calcium imaging to obtain a comprehensive, high-resolution spatial map of neural activity during ictal events. Wide-field calcium imaging confirmed the focal onset, with a sharply demarcated spread zone limited to 2-3 mm from the injection site (Figure 2). In some cases including the example shown, 4-AP was injected into visual rather than somatosensory cortex, indicating that the focality of 4-AP ictal events is regionally invariant.

**Figure 2.**
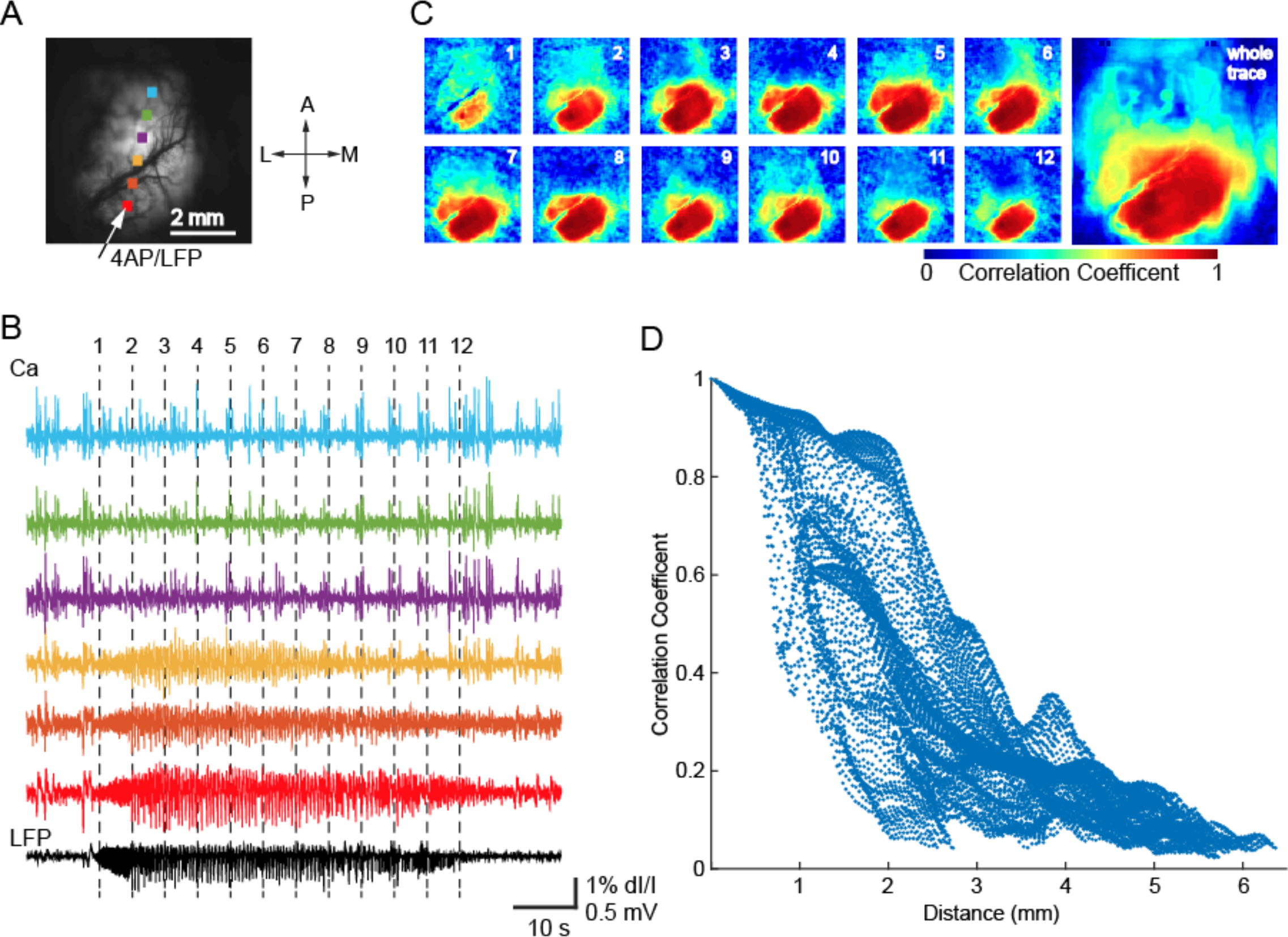
Wide field calcium imaging confirming 4AP ictal event focality. A. Field of view, with 4AP injection site and recording electrode indicated by the arrow. The dots show the locations of the corresponding color calcium traces in Panel B. B. LFP from the 4AP site (black, bottom trace) and calcium fluorescence (colored traces, 1 Hz high pass) during an ictal event. The vertical dotted lines indicate timing of the maps in Panel C. Note that the LFP pattern of the 4AP-induced ictal event is confined to the bottom 3 calcium traces, located within 2mm of the injection site. C. Spatiotemporal progression of the ictal event across the entire field of view. Left: seed-initiated correlation coefficient maps during the 12 time points marked in panel B. Note focal onset and limited spread of correlated calcium signal around the 4-AP electrode, supporting the focal nature of 4-AP ictal events. Right: seed-initiated correlation coefficient map of the entire ictal event. D. Graph of the Pearson correlation coefficient between each pixel’s calcium fluorescence and LFP amplitude plotted against distance of each pixel from the 4AP injection site. Note sharply decreased correlation values for distances greater than 2 mm.

### Multiunit activity at the ictal focus is associated with hemisphere-wide, distance-dependent LFP responses

The focal nature of 4-AP ictal events permitted investigation of neural responses at varying distances. During 4-AP ictal events as recorded with the microelectrode array, multiunit firing near the ictal center (≤1 mm) was associated with LFP fluctuations both locally and distally (Figure 3A). Spike-triggered averaging demonstrated distance dependence with respect to polarity and temporal delay (Figure 3B). Electrodes proximal to the ictal center (≤1*mm*) showed large negative LFP deflections that peaked shortly after multiunit firing bursts (Figure 3B, blue traces; median peak times: 4.3 ms, interquartile range: 3.5 to 5.4 ms; Figure 3C, blue arrows). Distally (> 4mm), there were positive deflections with prolonged delays (Figure 3B, green traces; median peak times: 17.6 ms, interquartile range: 14.9 to 18.9 ms; Figure 3C, red arrows). The distance-dependent polarity switch (Figure 3D, left subpanel, blue to red arrows) and temporal delay (Figure 3D, right subpanel, blue to red arrows) were observed across experiments (8 animals, 120 ictal events).

**Figure 3.**
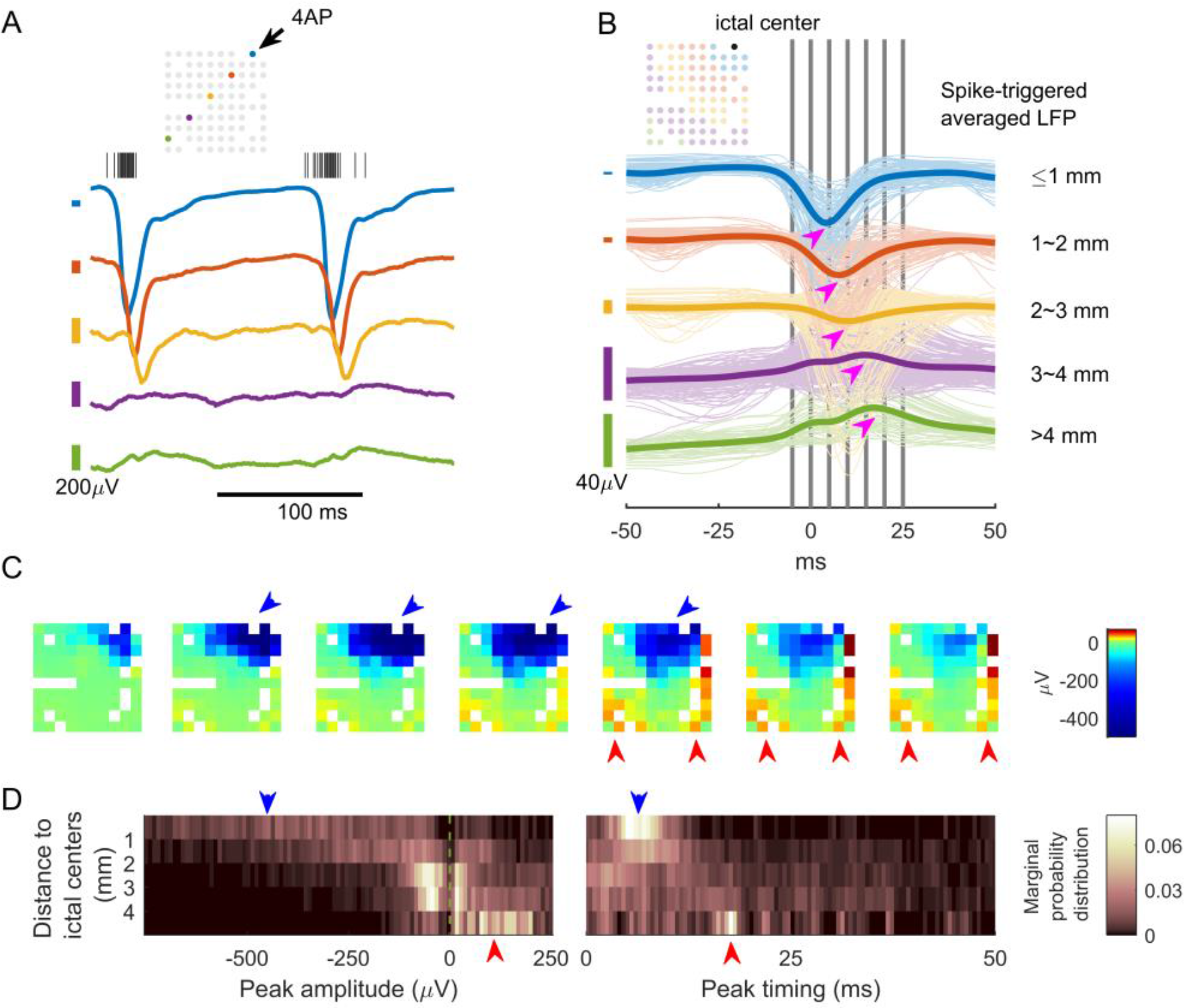
MUA at the ictal focus associated with hemisphere-scale, distance-dependent LFP responses. A. Sample LFP traces with ictal multiunit activity, color coded to indicate location of the recording microelectrode (inset schematic). For clarity of presentation, variable vertical scaling is used. The centers of the color-coded scale bars (left) are the isoelectric points for the corresponding trace. B. Spike-triggered LFP in one animal, averaged over 11 ictal events. Colors are used to indicate distance from the ictal center. The thick lines are the average of traces with similar colors. Notice the sequentially increased temporal delay of peak response and the switch of polarity (magenta arrows) as distance increases. Median delay for blue traces, measured from timing of the negative peaks: 4.3 ms (3.5 - 5.4 ms, n = 88). Median delay for green traces, measured from timing of the positive peaks: 17.6 ms (14.9 - 18.9 ms, n = 55). Delay between blue and green groups is different (U-test comparison of medians, p<<0.001). Vertical gray bars indicate the timing of the frames in Panel C. C. The spatiotemporal evolution of spike-triggered averaged LFP. Each frame’s time is indicated by Panel B’s vertical gray bars from left to right sequentially. Large negative LFP deflections were found near the ictal center (blue arrows) whereas positive LFP deflections were found distal to the ictal center, and with increased/different temporal delay (red arrows). D. Summary of spike-triggered averaging studies (8 animals, 120 ictal events). For each channel during each ictal event, the spike-triggered LFP’s global extremum (greatest deviation of signal from the isoelectric point) was recorded. The search range for global extrema was limited within the causal part of the signal (0 to +50 ms). For statistical analysis, data were grouped by anterior-to-posterior position with their extrema’s distribution shown in heat maps (5 groups, N = 1131, 1506, 2263, 1515, and 346 from proximal to distal). Left subpanel: distribution of extrema value, showing switch of polarity (probability of detecting a negative peak = 0.93, 0.75, 0.58, 0.45, 0.14, sign test,*α* = 0.05). Right subpanel: distribution of peak timing showing temporal delay (median: 6.9, 8.7, 10, 14.9, 17.5 ms, Kruskal–Wallis test, p<0.001).

### Parvalbumin(+) interneurons contribute to surround inhibition effect of ictal events

The observations of widespread, hemisphere-scale LFP effects and the proximal-distal polarity flip led us to hypothesize that ictal activities exert qualitatively opposite effects in distal compared to proximal neural tissues, and that the distal effect is inhibition-dominant (Prince and Wilder, 1967; Marshall *et al*., 2016). The greater distal temporal delay suggests that a multi-synaptic pathway is involved, consistent with feedforward inhibition. To provide support for our interpretation of the proximal-distal polarity flip, we employed in vivo 2-photon calcium imaging to record the activity of parvalbumin (PV)-positive interneurons in the distal surrounding tissue during ictal events (Figure 4A). We selected PV(+) interneurons because of their strong inhibitory effect on pyramidal neurons through proximal dendrite and soma projections (Markram *et al*., 2004; Kepecs and Fishell, 2014). Also, PV(+) interneurons have been reported to restrain ictal propagation in acute brain slices (Cammarota *et al*., 2013; Sessolo *et al*., 2015). Upon 4-AP injection at posterior primary visual cortex (V1), the ictal events reliably activated PV(+) interneurons in distal regions (imaged at ~4mm anterior to the 4-AP pipette tip in V1, Figure 4B-D). The recruitment of distal PV(+) interneurons to electrographic ictal events occurred with little temporal delay (Figure 4E-F, see legends for detailed statistics). The rapid response indicated that PV(+) interneurons are activated through synaptic transmission instead of slow processes such as extracellular ion diffusion. Overall, the observation of rapid PV(+) interneuron activation supported our hypothesis that ictal events exert strong, immediate inhibitory effects on distal tissues.

**Figure 4.**
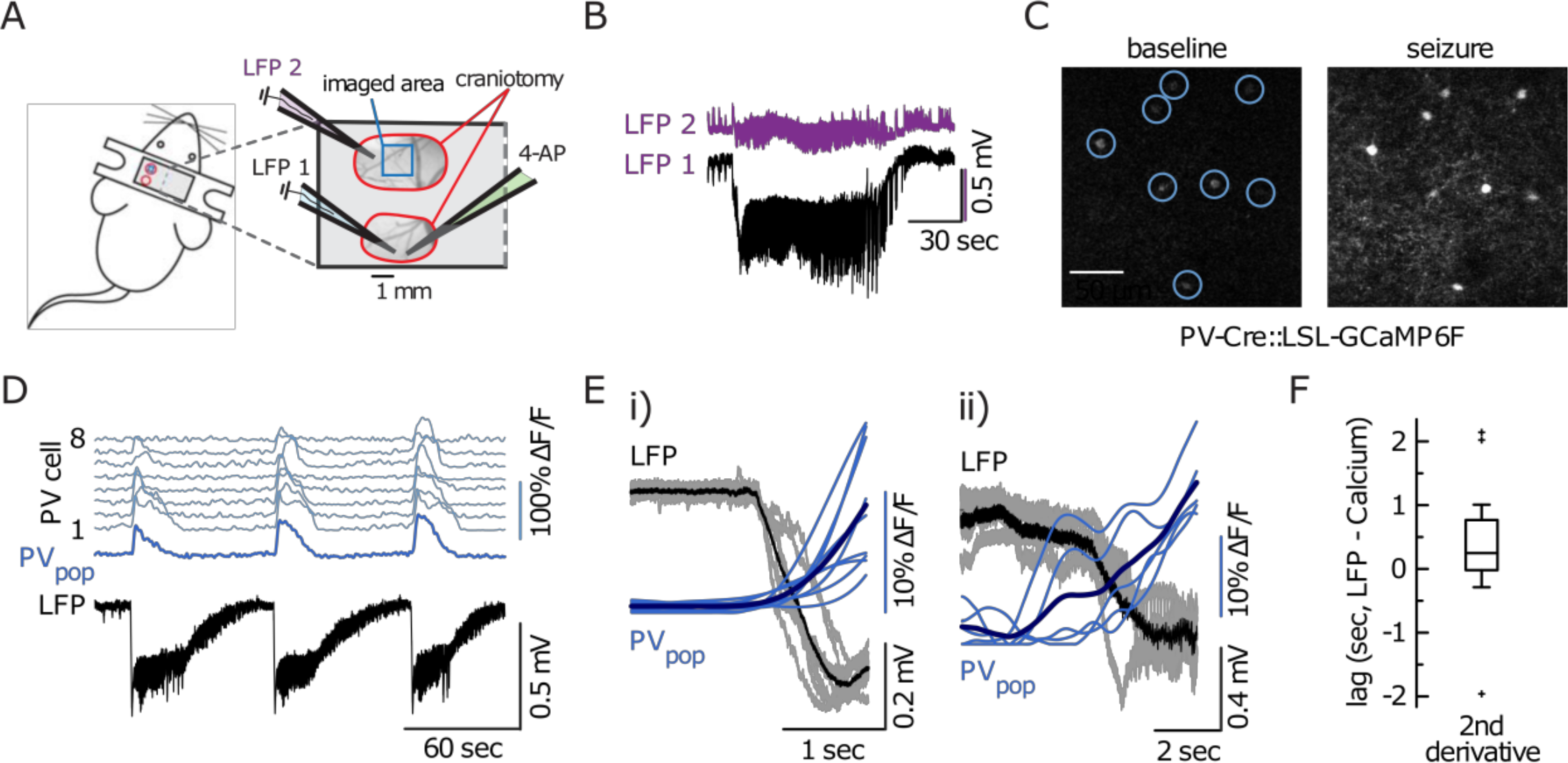
In vivo two-photon sub-population calcium imaging shows rapid recruitment of distal PV(+) interneurons to ictal events. A. Experimental setup: Two craniotomies over left primary somatosensory (SS1, just posterior to Bregma), and the posterior part of primary visual cortex (V1), respectively; each craniotomy is encircled in red; imaging field of view (FOV) outlined in blue. In addition to PV(+) subpopulation imaging, experiments involved the insertion of either two or three glass micropipettes: one pipette containing 4-AP in posterior V1 (green, 15mM, injection volume 500 nL), and one (injection site [LFP 1, light blue]) or two (injection site [LFP 1, light blue] plus imaging site ~4mm away [LFP 2, purple]) pipettes each containing a silver chloride silver for LFP recordings. B. Two simultaneous LFP recordings (LFP 2 [purple] close by the imaging area in SS1, LFP 1 [black] proximal to 4-AP injection site in V1). In line with experiments carried out in rats, there is a drastic reduction in amplitude and loss of DC-shift at the distal LFP electrode (inter-electrode distance ~4mm). C. Representative 3 second average calcium images depicting 8 distal PVs during baseline (left) and during ictal onset (right). The increase in calcium is visible to the naked eye. D. Calcium transients of the 8 PVs depicted in Panel C. Individual cells in light blue (top 8 traces), population average in dark blue. Simultaneous LFP recording near the 4AP injection site. Notice the consistent population recruitment across three consecutive ictal events. E. Superposition of all electrographic ictal onsets and corresponding PV population calcium transients (blue, mean in dark blue) in two independent experiments (i=9 ictal events, ii=5 ictal events). Note the systematic relationship between the paroxysmal DC-shift in the LFP (ictal onset) and a clear rise in the PV population calcium signal. F. Quantification of temporal PV population recruitment lag (distance to ictal initiation site ~4 mm). PV onset time points were defined by maximum 2nd derivative of the calcium signal within the peri-ictal onset time window (which was determined electrographically). Box plot box depicts 25 – 75 %ile, horizontal bar within box represents the median (248.47 msec, interquartile range 786.44 msec; 2 animals, 14 ictal events). Outlier values are displayed as +.

### Contiguously compromising GABA-A mediated fast inhibitory conductance causes classical ictal invasion

We subsequently tested the hypothesis that feedforward inhibition in the surrounding cortex is essential for preventing ictal propagation *in vivo*. We focused on pathways mediated by GABA-A conductance, because this is most consistent with the short temporal delay revealed by the spike-triggered LFP studies (Lopantsev and Avoli, 1998; Uva *et al*., 2005; Avoli and de Curtis, 2011; Pouille *et al*., 2013). After the 4AP injection was performed and 4-AP induced focal ictal events were observed, we bathed the whole cranial window with 5mM bicuculline (BMI, a GABA-A receptor antagonist) to achieve a contiguous breakdown of GABA-A mediated surround inhibition. This resulted in ictal activity propagating outward contiguously from the 4-AP site (3 animals, 22 ictal events), eventually invading the entire sampled area (Figure 5A-B). Notably, the morphology of the low-frequency LFP remained similar to the ictal events generated by 4-AP alone, as opposed to events triggered by focal BMI, which produces very high amplitude paroxysmal epileptiform discharges (Ma *et al*., 2004; Ma *et al*., 2009; Geneslaw *et al*., 2011). Multiunit activity showed parallel effects, with intense firing during ictal events at sites distant to the 4-AP injection site that were quiescent prior to BMI injection, and a visible recruitment wave with speeds increasing over time (Figure 5C-D). The ictal events also showed traveling waves originating from the 4-AP focus, as evidenced by sequential multiunit spike latencies (Figure 5E-F). In line with previous reports of ictal traveling waves in disinhibited rodent cortex (Chagnac-Amitai and Connors, 1989; Pinto *et al*., 2005; Trevelyan *et al*., 2007a), cross correlation of spike trains indicated average traveling wave speed of 63 mm/sec (Figure 5G, 95% confidence interval: 61.6 to 64.4 mm/sec). In summary, the morphological similarity of LFPs and the fact that ictal traveling waves solely originated from 4-AP injection sites suggested a single expanding ictal focus, analogous to the scenario during classical seizure propagation.

**Figure 5.**
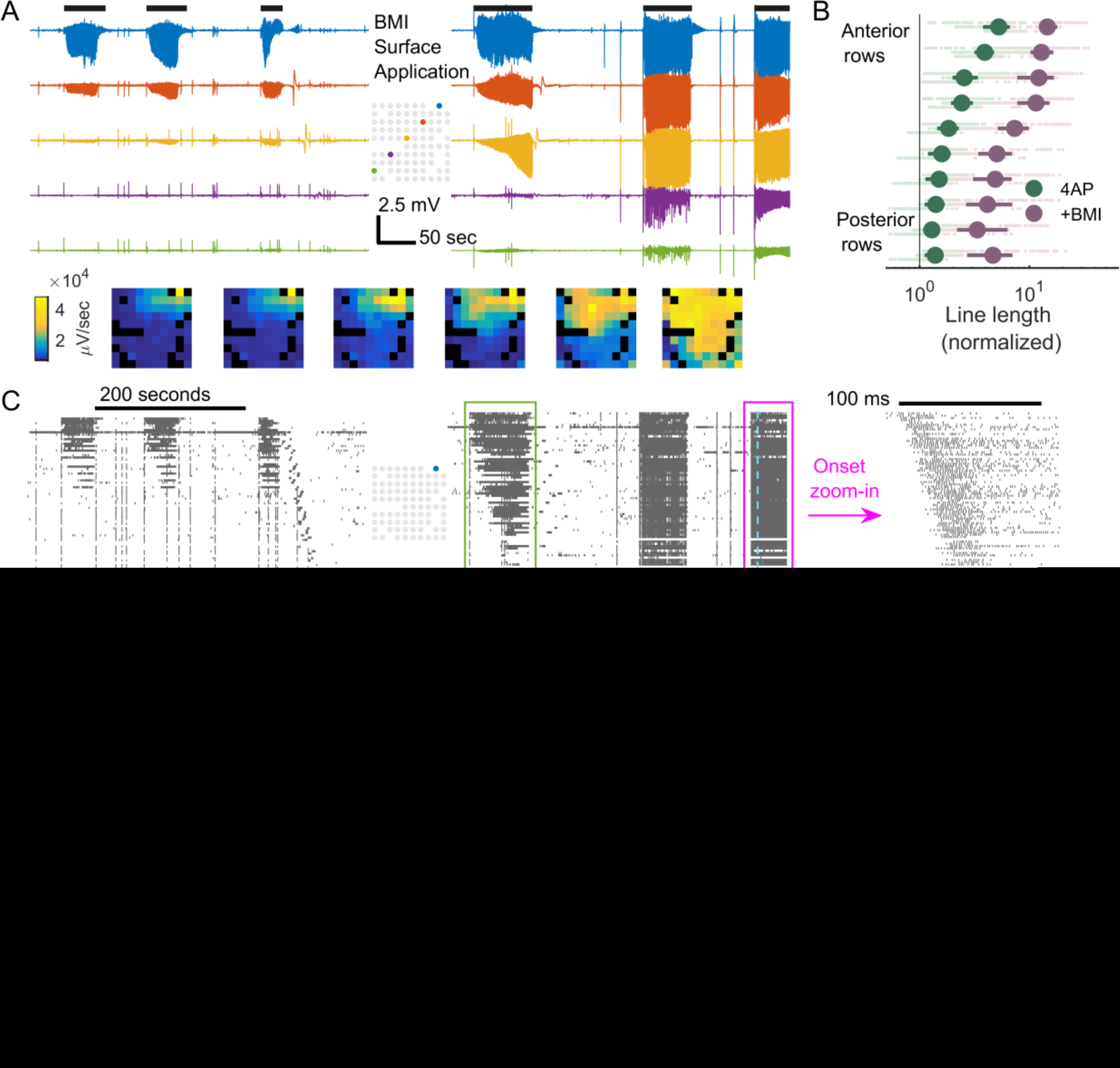
Contiguously reducing inhibition strength causes classical ictal propagation. A. LFP of selected channels before (left) and after (right) BMI surface application. Horizontal black bars (top) indicate ictal periods. The spatial distributions of the 6 ictal events, measured with line-length, are shown in frames below sequentially. B. Comparison of ictal activity (line length) before versus after BMI application. The panel’s conventions are analogous to Figure 1E. Each light green and red dot represents data from one channel during one ictal event. Spearman correlation coefficient between a channel’s position anterior-posteriorly and its line-length during ictal events after BMI application: 0.61 (3 animals, 22 ictal events, 1221 data points (electrode recording * ictal event), p << 0.001). C. MUA before (left) and after (right) BMI surface application. Time is aligned with Panel Data are sorted top to bottom according to distance from the ictal center (blue electrode of the inlet map). Green box: the first ictal event after BMI application. Notice the sequential recruitment of neurons from the 4-AP ictal center to distal areas. The ictal event marked with magenta box is zoomed in. Notice the onset of the ictal event with spatially sequential recruitment from the 4-AP injection site, albeit at higher speed than the one in the green box. D. The spatiotemporal evolution of a sample multiunit burst during an ictal event. The blue vertical bars indicate the timing of the frames (right). Notice the spread of the traveling wave pattern radiating from the 4-AP ictal center. E. Cross-correlogram of the spike train recorded from the 4-AP ictal center to other channels during all ictal events (5 events) after BMI application. The colors indicate the distance between the channel’s distance from the 4-AP ictal center. F. Pairwise cross-correlation of the same dataset. Channels are sorted horizontally and vertically by distance from the 4AP ictal center. Times of peak cross-correlation are indicated by color. Only channels with average multiunit firing rate > 5 spike/s are included (66 channels). G. Linear regression of data from Panel H. Linear regression model : *d_i_ – d_j_ = V \** (*t_i_*–*t_j_*) where *i* & *j* are channel indices, *d* is the distance from the 4AP ictal center, and *V* is speed. Least squares regression shows V = 63 mm/sec, F-test p << 0.001, adjusted *R*^2^ = 0.648 (66 channels, 2145 pairs).

### Long-range ictal propagation with dual foci

The observations of distant (> 4mm) LFP signal (Figure 1B, 1E, Figure 3) as well as triggered PV(+) interneuron activity (Figure 4) suggested a hidden, large-scale excitation pathway. We therefore predicted that ictal propagation to a distant focus could occur due to locally impaired inhibition at the distant site. To induce this condition, a solitary focal BMI microinjection (5 mM, 500 nL) was placed at a distant, non-contiguous region 4-5 mm from the 4-AP focus, with both foci in the same hemisphere. This procedure resulted in ictal activity involving both the 4AP and BMI injection sites (N=5 animals and 45 ictal events, Figure 6A, top). In contrast to the classical propagation (Figure 5), the strongest multiunit activation was demonstrated at the two microinjected sites with nearly simultaneous onset (Figure 6A, bottom). Moreover, there was markedly different temporal evolution demonstrated in both LFP and multiunit activity (Figure 6A-B, Supplementary Figure S2). In two animals, we were able to discern an area with initially low multiunit firing rates in the region between the two foci, further indicative of non-adjacent seizure propagation via a large-scale excitatory pathway (Figure 6B-C). This in-between area was recruited antidromically (Figure 6C, posterior to anterior, average speed: 0.19 mm/sec, 95% confidence interval: 0.17 to 0.21 mm/sec, 21 ictal events). The antidromic recruitment indicated that the BMI focus, even though it should not have been able to generate a seizure resembling the 4-AP events independently, acted as a secondary driver of ictal activity instead of simply mirroring projections from the 4-AP ictal focus.

**Figure 6.**
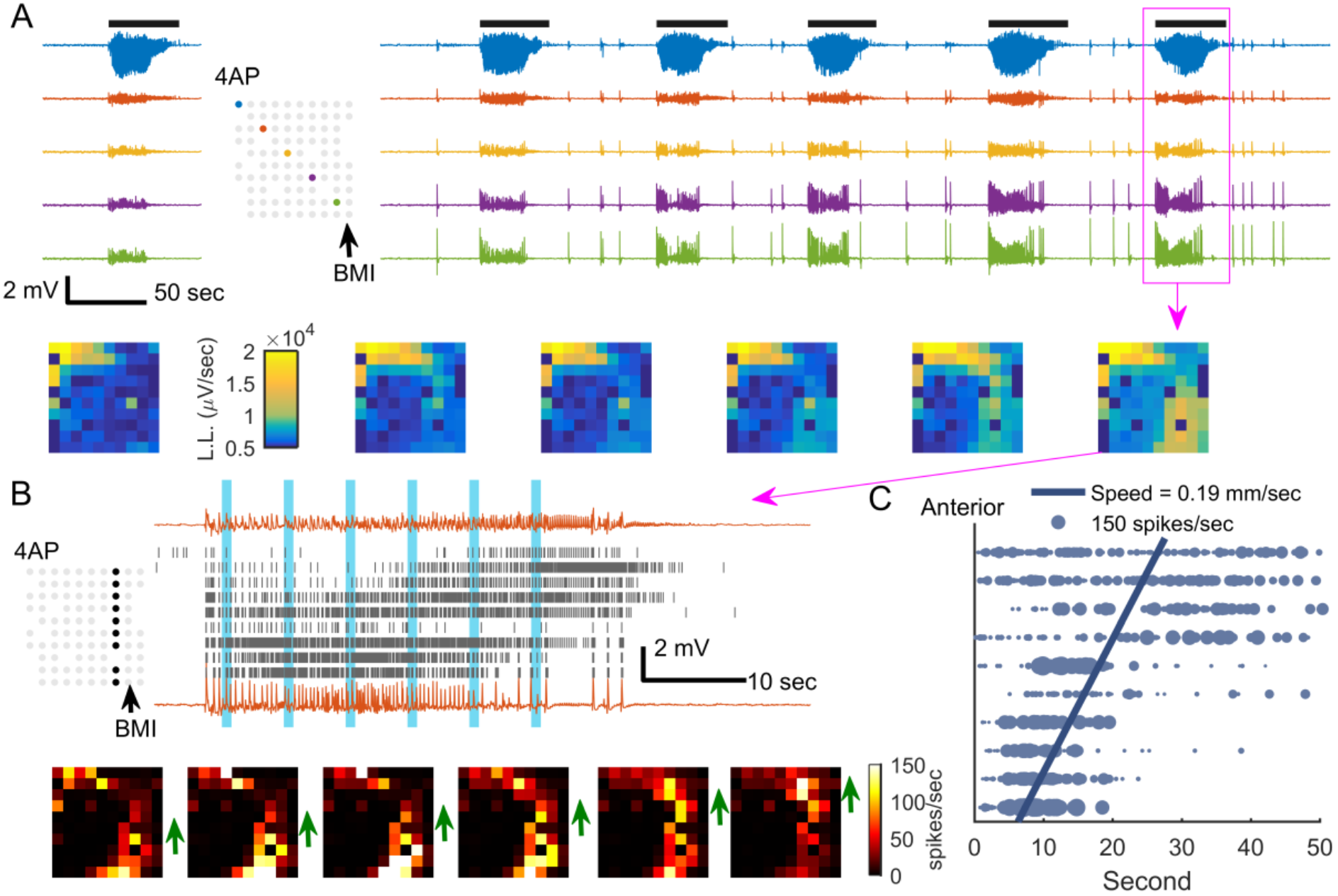
Emergence of two ictal foci through long-range ictal propagation pathways. A. Raw LFP traces before (left) and after (right) BMI injection. Figure conventions are adopted from Figure 5A. Note that the ictal event at the BMI site is temporally linked with the 4-AP event but with a quite different electrographic morphology (blue vs. green traces; see higher resolution image in Supplementary Figure S2). Frames (bottom) show the spatial distribution of each ictal event sequentially. L.L.: line-length. B. MUA during the ictal event marked with a magenta box in Panel A. Each row of the raster plot was recorded from the electrode whose physical position is marked in the inset map (upper: anterior electrodes; lower: posterior electrodes). The upper and lower traces are the LFP recorded from the most anterior and the most posterior electrodes of the marked electrode column respectively. Both the raster plot (top) and snapshots of the entire microelectrode array (bottom) demonstrate the slow progression of sustained, intense multiunit firing rates (1s Gaussian kernel) that characterizes ictal recruitment from the location of the BMI focus (arrow), toward the area near the 4-AP focus (green arrows). The timing of each snapshot corresponds to the vertical blue bars in the raster plot. We refer to this as antidromic recruitment, as the bottom to top progression is the reverse of what would be expected with classical seizure propagation. C. Statistical analysis of anterior-posterior propagation of 4-AP ictal events after BMI injection. Horizontal coordinates indicate timing relative to event onset of the maximal firing rate (1 s Gaussian kernel), and vertical coordinates (*x*) indicate channel rows (anterior-posterior as in prior figures). Each datum was plotted with marker size proportional to its maximum firing rate. Weighted linear regression model: *t = β _0_* + *β*_1_*x*. 846 data points, 21 ictal events, 95% confidence interval: 0.17 to 0.21 mm/sec, F-test, p<<0.001, adjusted *R*^2^ = 0.343.

We next sought to replicate the creation of a dual-focus network across longer distances using wide field calcium imaging with similar experimental methods in 8 animals. In 7 of the animals, two linked foci at the two injection sites were seen (Figure 7). In all cases, ictal events with morphology resembling 4AP seizures initiated approximately simultaneously at both injection sites (Figure 7C). However, calcium dynamics showed strong synchrony with the immediately adjacent area corresponding with a 2-3 mm focus at each site, with relatively weaker synchronization across the foci (Figure 7D).

**Figure 7.**
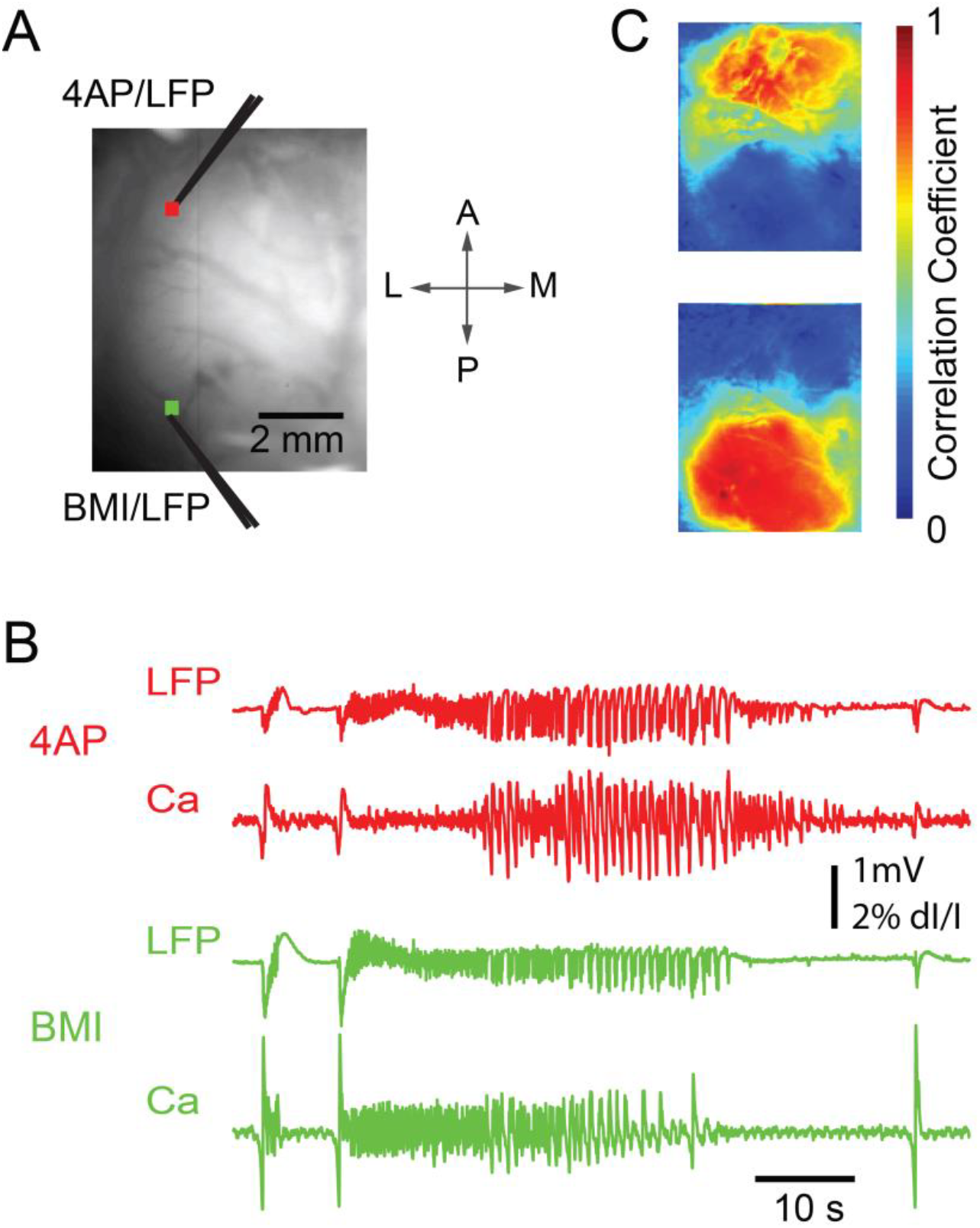
Wide-field calcium imaging of the two-focus epileptic network. A. Wide field imaging setup of the two-focus experiment. The cranial window extended from ~1mm caudal to bregma to ~1mm rostral to lambda, and was placed 1-6 mm from the midline. Distance between foci varied from 3.8 to 6.3 mm (mean 5.1 mm). The field of view shows the location of 4-AP and BMI injection sites. Glass electrodes were used for both microinjection and LFP recording. Two 3 × 3 pixel ROIs were selected close to the 4-AP injection site (red) and BMI injection site (green), from which the calcium trace was drawn in B) and selected as the seed trace for cross-correlation. B. LFP (top) and Ca signals (bottom) recorded from the 4-AP (red, top) and BMI (green, bottom) injection sites, respectively. Note that although the ictal events begin simultaneously, the evolution of morphology of the ictal waveform is quite different as recorded both with the LFP and the calcium imaging. C. Seed-initiated correlation coefficient maps (seed traces shown in panel B), with correlation studied for the entire set of 8 rats and 41 seizures. Top: Correlation map of a sample seizure in which the Ca signal recorded from the 4-AP injection site ROI was used as the seed trace. Mean and SD of the correlation coefficients calculated from a 1 mm × 1mm brain region surrounding the 4-AP injection site was 0.83 +/- 0.15, and for BMI site 0.31 +/- 0.21. The difference was highly significant (Wilcoxon rank-sum test, p<<0.001). Bottom: Correlation map of the same seizure, using the Ca signal recorded from the BMI injection site ROI as the seed trace. Correlation for the BMI site was 0.92 +/- 0.08, and for the 4-AP site 0.34 +/- 0.23, again significantly different (Wilcoxon rank-sum test, p<<0.001).

### Failure of dual focus recruitment via cross-callosal projections

Finally, we investigated whether the same network dynamics could be served by cross-callosal synaptic projections, by creating an area of focal disinhibition in the contralateral hemisphere. A 4-AP ictal focus was created in left somatosensory cortex, and BMI was injected in the contralateral (right) homotopic region. As expected, 4-AP ictal events emerged with field restricted to the ipsilateral hemisphere, while focal independent interictal spikes were observed at the contralateral site (Figure 8A). Ictal events did not propagate to the contralateral hemisphere, with no difference in contralateral LFP activity between ictal and interictal periods (Figure 8B) (p=0.805, 2 animals, 17 ictal events, 680 data points). Similarly, there was no consistent increase in average multiunit firing rates in the contralateral cortices during ictal periods in comparison to interictal periods (p = 0.98, Figure 8C-D). This finding was replicated in wide field calcium imaging (Figure 8E-F). Calcium fluorescence dynamics in the contralateral hemisphere showed little correlation to the 4-AP ictal focus (Pearson’s correlation coefficients: 0.068 ± 0.229, n = 7 animals). These results indicate that the likelihood of recruiting and synchronizing a second epileptic focus via cross-callosal projections is reduced in the acute rodent model (Figure 8F).

**Figure 8.**
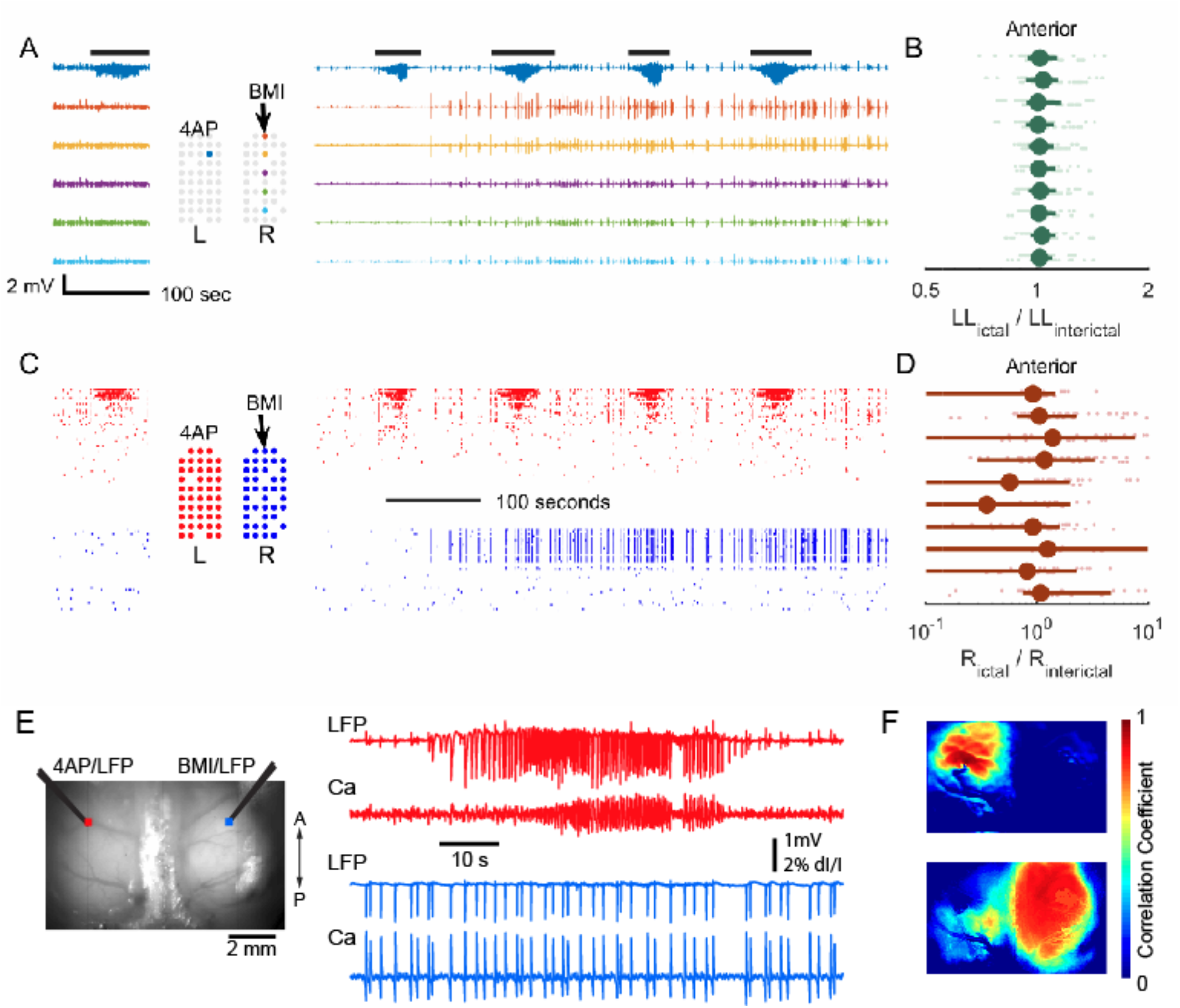
Contralateral BMI injection failed to induce cross-hemisphere propagation. A. 4-AP is injected to create a seizure focus in the left hemisphere. LFP before (left) and after (right) BMI injection in the contralateral homotopic somatosensory cortex. After BMI injection, paroxysmal interictal spikes were seen in the LFP; however, a second ictal focus did not develop. B. Ictal line-length relative to the interictal period following each ictal event. Data were plotted similarly to Figure 1E. Median 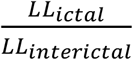=1.02, 17 ictal events from 2 animals, 675 data points, sign test p=0.805. C. Multiunit raster plot before (left) and after (right) BMI injection. Time is aligned with Panel A. D. Average ictal multiunit firing rate (R) relative to interictal, analogous to Panel B. Median 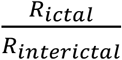=1.02, sign test, p=0.98. E. Wide field calcium imaging of 4-AP ictal focus in the left hemisphere and focal disinhibition with BMI injected at the contralateral homotopic cortex. Left: the field of view showing the injection sites. Right: The LFP and calcium fluorescence signal recorded from 4AP (top) and BMI (bottom) injection sites. The ictal events do not propagate to the contralateral side and the BMI injection results in focal interictal spikes without creating a new interconnected ictal focus. F. Seed-initiated correlation coefficient maps. Top: the seed trace used was taken from an ROI adjacent to the 4-AP injection site. Bottom: the seed trace used was taken from an ROI adjacent to the BMI injection site. The ictal event remains focal and does not involve the other hemisphere. The disinhibited interictal spiking zone is widespread and completely independent from the seizure focus.

## Discussion

We investigated local and distant neural responses to focal neocortical ictal events in an acute pharmacologically induced seizure model. Ictal events triggered by focal application of 4-AP remained confined to a small cortical territory, but displayed various propagation patterns depending on the site of BMI application. Two distinct propagation patterns were seen. Global BMI application resulted in a propagating ictal focus with a single dominant dynamic, whereas distant focal application resulted in the emergence of a second, non-synchronous ictal core. The observed spread patterns and temporal delays of ictal discharges were consistent with activation of large-scale excitatory synaptic pathways, with the success of seizure invasion depending on impairment of the GABA-A mediated inhibitory restraint. These data illustrate a scenario in which large-scale seizure spread and epileptic network formation may coexist with the classical model of contiguous cortical Jacksonian propagation, both involving the breakdown of inhibitory restraint. While the coupling of the two foci occurred acutely, in the absence of pathologically reinforced structural alterations, such alterations could enhance this effect.

Previously, we characterized brain areas where principle neurons demonstrate intense, synchronized burst firing (i.e. clonic firing) temporally locked to LFP discharges as the “ictal core” (Schevon *et al*., 2012). This region is sharply demarcated and significantly smaller than the area where large amplitude epileptiform LFP discharges can be recorded (Schevon *et al*., 2012; Smith *et al*., 2016). Regions outside the ictal core where epileptiform LFPs can still be observed but its neurons are not directly participating in the seizure are termed the “penumbra”. Synaptic activity in the penumbra is time-locked to neuronal activities in the ictal core, consisting of a mixture of excitatory and inhibitory field potentials, with pyramidal cell firing restrained by the inhibitory veto. This results in relatively sparse, heterogeneous firing that is generally not phase-locked to the low-frequency LFP (Truccolo *et al*., 2011; Schevon *et al*., 2012; Weiss *et al*., 2013). Thus, in the penumbra, the effects of strong widespread excitation are masked by local inhibitory responses. In our acute rodent neocortical model, this brain region extends far beyond the immediate surround, as evidenced by the fact that seizures can propagate rapidly to distant sites once the inhibitory veto is lifted.

The core-penumbra coupling mechanism is suggested in the morphology of LFP deflections associated with synchronized neuronal bursts during ictal events. The observed polarity flip (proximal-negative versus distal-positive) and temporal delay (~10 ms) suggests that nearby regions are dominated by monosynaptic local recurrent excitatory projections arising from pyramidal neurons within the ictal zone, while distant territories are dominated by multisynaptic feedforward inhibition, involving projections to inhibitory interneurons (Marshall *et al*., 2016). These observations are not consistent with a passive volume-conducted process, as we would then expect preserved polarity and minimal temporal delays. Interestingly, a similar distance-dependent polarity flip has been detected in human seizure recordings (Eissa *et al*., 2017).

We selected the 4-AP model (Szente and Pongracz, 1979; Szente and Baranyi, 1987; Ma *et al*., 2009; Zhao *et al*., 2009; Zhao *et al*., 2011; Ma *et al*., 2013; Zhao *et al*., 2015; Wenzel *et al*., 2017) due to its ability to trigger a well-defined, spatially limited (~2-3 mm diameter) neocortical ictal event, electrographically mimicking spontaneous human neocortical seizures with a low-voltage fast activity onset pattern. We demonstrated that the region of active spiking remains focal throughout the evolution of the seizure, as long as the surrounding inhibitory mechanisms are intact. This focality permitted distant BMI site placement in order to assess interactions between the core and surrounding areas. In a chronic model, the site of seizure origination is less well defined. While our use of an acute anesthetized model is a potential study limitation, it was necessary in order to create the dual-focus structure needed to test our hypothesis. Although the use of anesthesia may alter global excitation levels in comparison to the unanesthetized wake or sleep state, we have shown preservation of focal seizure spread dynamics in our anesthetized model, with electrographic characteristics resembling that seen in an unanesthetized whole-brain preparation (Uva *et al*., 2013). Focal, spreading seizures are also known to occur from deep anesthesia in the clinical setting (Howe *et al*., 2016). The use of an acute model also allowed us to determine that complex epileptic network interactions can occur in the absence of pathologically reinforced connectivity. Such pathological high connectivity may, however, enhance the likelihood of distant recruitment. For example, the failure of cross-callosal recruitment in our model does not imply that this cannot occur in human epilepsy.

We found evidence that the two distinct seizure propagation patterns can both be explained by inhibition breakdown. At the juxta-ictal core region, both synaptic as well as extracellular mechanisms, i.e. potassium diffusion and field effects, may contribute to the breakdown of inhibition (Pumain *et al*., 1985; Bikson *et al*., 2003; Park and Durand, 2006; Frohlich *et al*., 2008; Mylvaganam *et al*., 2014; Zhang *et al*., 2014), resulting in classical Jacksonian march-like propagation (Schevon *et al*., 2012; Wenzel *et al*., 2017). In contrast, inhibition compromise and seizure emergence at a distal site may be attributed to synaptic or circuit mechanisms, such as intracellular chloride accumulation (Huberfeld *et al*., 2007; Barmashenko *et al*., 2011; Lillis *et al*., 2012; Buchin *et al*., 2016), synchronization (Lehnertz *et al*., 2009; Jiruska *et al*., 2013), or regional variation of inhibition robustness. This is further supported by a recent study of a picrotoxin mouse model in vivo, in which visual cortex seizures showed either local, lateral propagation or homotopic spread to functionally connected sites (Rossi *et al*., 2016). Two dynamically distinct ictal foci emerged in the experiments involving separate 4AP and BMI injection sites. In some instances, seizure activity was restricted to regions close to the two foci, with the seizure appearing to “jump” across the gap. A key observation regarding the epileptic activity observed at the BMI site is that the oscillatory pattern was similar to 4-AP triggered seizures, rather than the high amplitude bursts of the BMI model, demonstrated in the contralateral BMI injection experiment (Figure 8). The appearance of a 4AP-type seizure at the BMI injection site is thus a strong demonstration that the source of the seizure at this site was the distant 4-AP focus, rather than being solely attributable to BMI.

The nearly simultaneous ictal event onsets at the 4-AP and BMI foci, consistent with the rapid activation (on a scale of milliseconds) of distant interneuronal activity seen in the two-photon imaging data, would be hard to tease apart by examining LFP or EEG data alone. This would be especially true if there is a variable relationship of the LFP recording site to the location of seizure emergence, as is generally the case in clinical recordings. This may also occur in chronic experimentally-induced seizure models, due to uncertainty regarding the precise location of spontaneous seizure emergence.

These experiments provide proof of principle that seizures can generate secondary foci that appear to operate as a unified network, when viewed in wideband recordings. If this same process occurs in humans, there are immediate implications for clinical treatments that target seizure foci, such as epilepsy surgery. The originating seizure focus may induce scattered dependent foci in outlying brain areas that may or may not be capable of triggering a seizure independently, but may manifest immediately upon seizure onset. These may obscure the location and extent of the originating focus, with attendant consequences for surgical decisions. In addition, the same process studied here may operate across seizure foci that are capable of triggering seizures independently, although mechanisms apart from depressed local inhibition may contribute to the formation of such foci.

In conclusion, our study shows that focal ictal events project strong and widely distributed excitation in an acute rodent seizure model. Compromised local inhibitory response, depending on its distance from the ictal core, can result in various seizure propagation patterns and epileptic network formation. Further studies to confirm these findings and investigate the various distance-dependent seizure defense mechanisms involved should be conducted using different acute and chronic seizure models, including in unanesthetized animals, as well as alternative methods of impairing inhibition and perhaps focusing on different interneuronal populations. Additionally, further studies to provide direct evidence of this process in humans will help to establish its potential role in informing clinical management and development of new treatments.

## Acknowledgements

We acknowledge the useful discussions with Dr. LF Abbott, Dr. Kenneth Miller, and Dr. Steven A Siegelbaum. H.M. and T.S. acknowledge funding from CURE and the National Science Foundation (NSF-1264948); J.L. and C.A.S. acknowledge funding from the Simons Foundation and NIH/NINDS R01 NS084142. R.Y.’s laboratory is supported by the NEI (DP1EY024503, R01EY011787), NIMH (R01MH101218, R01 MH100561). This material is based upon work supported by, or in part by, the U. S. Army Research Laboratory and the U. S. Army Research Office under contract number W911NF-12-1-0594 (MURI).

## Author contributions

CAS, RE, and THS conceived of the project and were directly involved in initial animal studies (Utah array recordings) at Cornell. JYL, HM, MZ, AD, and EBD conducted the animal experiments at Cornell (Utah array and calcium imaging) and performed the bulk of the data analysis. JYL, EHS, CAS, RE, and THS participated in data interpretation and analysis. The two-photon imaging of parvalbumin+ interneurons was conceived by JYL and conducted by MW and RY.

---

**Figure S1.**
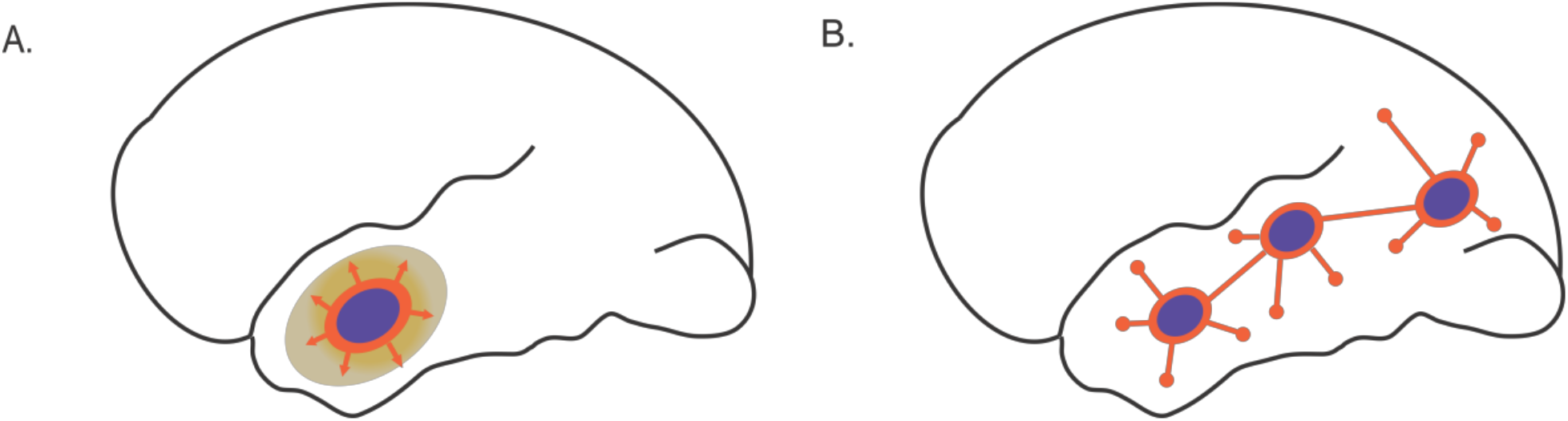
Two topological patterns of focal seizure propagation. A. Classical, Jacksonian-like focal seizure propagation. The purple zone is the ictal core, surrounded by the ictal wavefront (orange), slowly expanding toward adjacent cortical circuitry. The yellowish area surrounding the ictal core represents the ‘ictal penumbra’, an area that receives massive synaptic barrages from the ictal core and demonstrates epileptiform EEGs but has not been recruited into the seizing territory. B. The epileptic network. Several disparate ictogenic zones (purple) are recurrently connected by synaptic projections (orange lines). Focal seizure activity may arise from coordinated activity and rapidly propagate to distant and disparate regions through distributed mechanisms.

**Figure 2.**
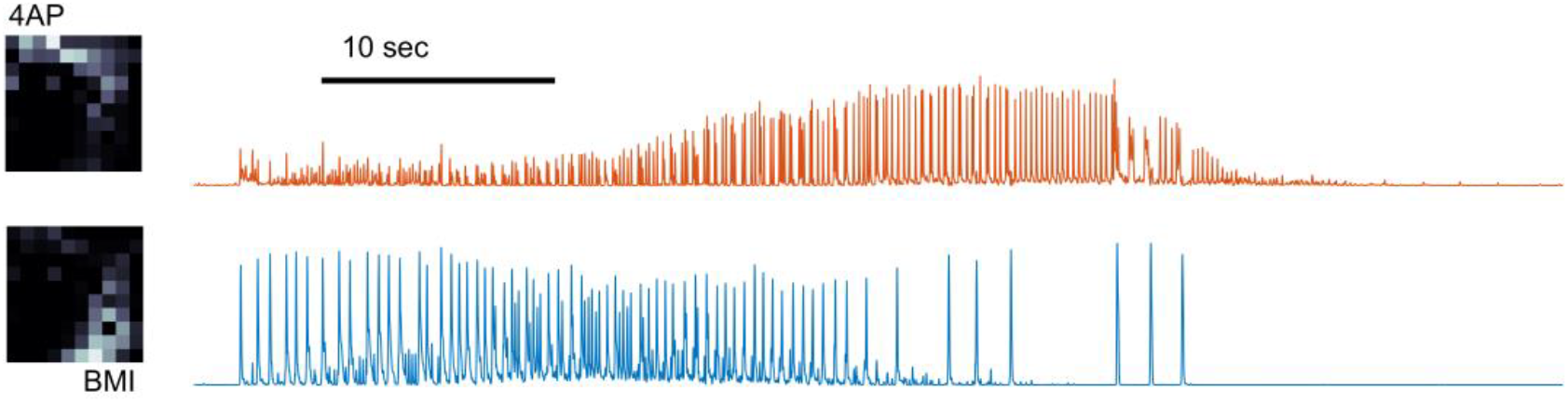
Multiunit dynamics analysis under non-contiguous inhibition breakdown shows complex interactions between the two ictal foci. Spatiotemporal dynamics of multiunit activity during a sample ictal event, analyzed by non-negative matrix factorization (see methods: cluster analysis of multiunit spike trains, residual Frobenius norm divided by the original matrix norm: 44%). Left panels: the spatial distribution of the two non-negative factors. The sites of maximal activity spatially correspond to the two microinjection sites (text labels). LFP traces from the channel with maximal multiunit firing rate at each site (4-AP = orange, BMI = blue) demonstrate the contrasting temporal dynamics of the dual ictal foci. Note that the ictal-associated activity begins roughly simultaneously at both sites, but the waveform morphologies evolve independently and terminate at different times. The BMI site trace (blue) demonstrates initial bursting that is characteristically induced by this agent, but this evolves into a morphological pattern characteristic of seizures induced by 4-AP.

